# Functional connectivity in neuromuscular system underlying bimanual muscle synergies

**DOI:** 10.1101/056671

**Authors:** Ingmar E. J. de Vries, Andreas Daffertshofer, Dick F. Stegeman, Tjeerd W. Boonstra

## Abstract

Neural synchrony has been suggested as mechanism for integrating distributed sensorimotor systems involved in coordinated movement. To test the role of corticomuscular and intermuscular coherence in the formation of bimanual muscle synergies, we experimentally manipulated the degree of coordination between hand muscles by varying the sensitivity of the visual feedback to differences in bilateral force. In 16 healthy participants, cortical activity was measured using 64-channel electroencephalography (EEG) and muscle activity of the flexor pollicis brevis muscle of both hands using 8×8-channel high-density electromyography (HDsEMG). Using the uncontrolled manifold framework, coordination between bilateral forces was quantified by the synergy index *R_V_* in the time and frequency domain. Functional connectivity was assed using corticomuscular coherence between muscle activity and cortical source activity and intermuscular coherence between bilateral EMG activity. As expected, bimanual synergies were stronger in the high coordination condition. *R_V_* was higher in the high coordination condition in frequencies between 0 and 0.5 Hz, and above 2 Hz. For the 0.5-2 Hz frequency band this pattern was inverted. Corticomuscular coherence in the beta band (16-30 Hz) was maximal in the contralateral motor cortex and was reduced in the high coordination condition. In contrast, intermuscular coherence was observed at 5-12 Hz and increased with bimanual coordination. Within-subject comparisons revealed a negative correlation between *R_V_* and corticomuscular coherence and a positive correlation between *R_V_* and intermuscular coherence. Our findings suggest two distinct neural pathways: (1) Corticomuscular coherence reflects direct corticospinal projections involved in controlling individual muscles; (2) intermuscular coherence reflects diverging pathways involved in the coordination of multiple muscles.

## 1. Introduction

The many degrees of freedom (DOFs) of the musculoskeletal system provide great flexibility, but they make the corresponding control problem rather complex. There are numerous ways to perform a movement to achieve the same goal. How does the nervous system coordinate the abundant DOFs? To reduce the dimensionality of the control problem, it has been proposed that a group of muscles spanning several different joints can become functionally linked so as to behave as a single task-specific unit (Bernstein 1967; Latash et al. 2007; Turvey 1990). The nervous system can control these muscle synergies - rather than individual muscles - and by combining muscle synergies in different ways, a small number of synergies can account for a wide variety of movements (d’Avella et al. 2003; Ivanenko et al. 2004; Ting and Macpherson 2005; Tresch et al. 1999). However, relatively little is known about how these muscle synergies are implemented in the central nervous system (Tresch and Jarc 2009). The spinal circuitry and the pathways connecting the motor cortex and the spinal interneurons appear central in understanding the neural basis of muscle synergies (Bizzi and Cheung 2013; Levine et al. 2014).

Neural synchronization may offer a window into the neural correlates of muscle synergies (Boonstra 2013). Equivalent to its role in perceptual processes (Engel et al. 2001; Singer 1999), neural synchrony may provide a mechanism for integrating the distributed motor and sensory systems involved in coordinated movement and posture (Farmer 1998). If neural synchrony is a general mechanism for coordinating neural activity, one might expect that muscles taking part in a synergy receive correlated neural input. In fact, neural synchrony has been widely observed in the motor system (Schnitzler and Gross 2005; van Wijk et al. 2012). In particular, corticomuscular coherence has been observed between motor cortex and muscle activity (Baker et al. 1997; Conway et al. 1995; Salenius et al. 1997), and intermuscular coherence between different sets of muscles (Boonstra et al. 2008; Farmer et al. 1993; Kilner et al. 1999). Interestingly, muscles contributing to a synergy generally reveal intermuscular coherence suggesting they receive correlated neural input (Boonstra et al. 2009a; Danna-Dos-Santos et al. 2014; De Luca and Erim 2002; Laine et al. 2015).

In this study we tested the relationship between muscle synergies and intermuscular and corticomuscular coherence by experimentally manipulating the required level of
coordination between bilateral hand muscles. Participants performed a bimanual precision-grip task while EEG and EMG activity were recorded (cf. Boonstra et al. 2009b). Similarly to Nazarpour et al. (2012), we controlled the degree of bimanual muscle synergy through visual feedback presented to the participants by varying the sensitivity of the to-be-controlled cursor to differences in the two hand forces, making this difference the relevant dimension for task success, while making the sum of the two hand forces the (relatively) irrelevant dimension for task success. That way we could experimentally manipulate the amount of variability in the relevant versus the irrelevant dimension for task success (cf. Scholz and Schoner 1999).

We expected to find higher levels of intermuscular coherence between homologous hand muscles but lower levels of corticomuscular coherence when a stronger synergy is required. Corticomuscular is thought to reflect direct corticospinal projections (Williams and Baker 2009), involved in the selective control of individual muscles (Lemon 2008). We hence expect that corticomuscular coherence is more dominant when no bilateral coordination is required and both hands can be controlled independently. This would imply that our nervous system can flexibly regulate the contribution of muscle synergies depending on task demands and that the neural basis of muscle synergies can be assessed using functional connectivity analysis between muscle and cortical activities. The level of bilateral muscle synergy was quantified from the force trajectories using a measure of synergy proposed by Latash et al. (2002). We extended this measure to the frequency domain to investigate the time scale at which the bilateral synergy is established. Establishing a link between functional connectivity and muscle synergies will help to delineate the neural pathways involved in motor coordination.

## 2. Materials and Methods

### 2.1 Participants

Sixteen healthy right-handed participants (8 female, age = 25.5 ± 4.3) volunteered in the study. The local ethics committee at the Faculty of Human Movement Sciences, VU University Amsterdam, approved the experimental protocol and each participant provided written informed consent prior to participating in the experiment.

### 2.2 Experimental protocol

Participants performed a bimanual precision-grip task while EEG and EMG activity was recorded. Bimanual forces were generated against two compliant force sensors held in each hand using a pinch grip (Fig. 1A).

**Figure 1.**
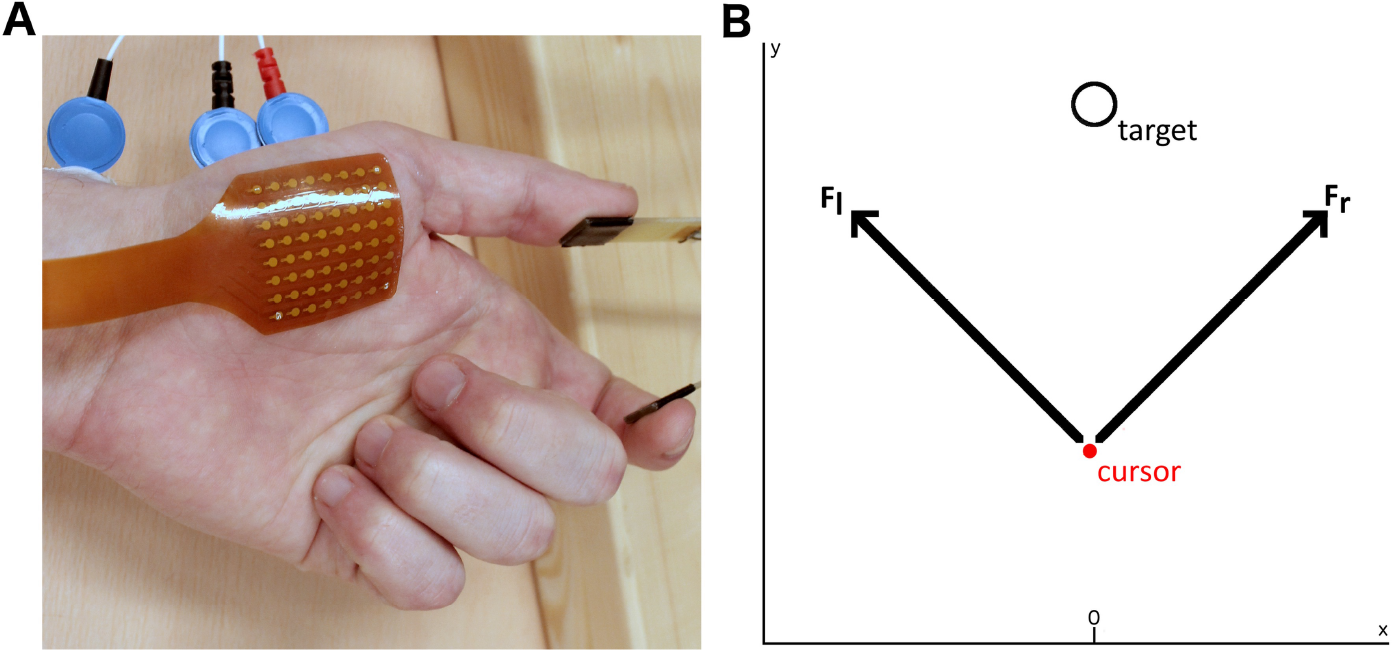
Experimental protocol. **A)** High-density electrode grid is placed over the flexor pollicis brevis while participants perform a pinch grip on custom-made compliant force sensor **B)** Force level projection. Cursor and target are the visual feedback displayed on the monitor.

Participants were instructed to track a visual target by moving the cursor displayed on a monitor (Fig. 1B). At the start of each trial, the target was in the starting position. During the first 5s, the target moved linearly to the final target position (force ramp), where it stayed for the remaining 10s of the trial (constant force). Participants could move the cursor by applying force to both sensors in order to keep the cursor within the target. The position of the cursor was determined by a linear weighting of the forces generated by the participant:

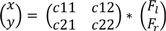

where *x* and *y* are the horizontal and vertical screen coordinates and *F_l_* and *F_r_* are the force levels generated by the left and right hand, respectively (Fig. 1B). The 2×2 projection matrix *c* converts the generated force levels to screen coordinates. The required level of bilateral muscle synergy was experimentally manipulated by changing the projection matrix *c* (Nazarpour et al. 2012). For example, when

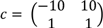

the *x*-position of the cursor is relatively sensitive to differences between the two force levels so a relatively strong coordination between both hands is required to perform the task successfully (i.e. keep the cursor in the circular target). That is, even a small difference between *F_l_* and *F_r_* will move the cursor sideways. In contrast, when

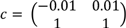

the *x*-position of the cursor is relatively insensitive to differences between the two force levels, i.e. a relatively weak coordination between both hands is required. For the latter, the total force needs to be controlled to move the cursor up or down, but a large variability in the ratio of the two force levels is tolerated (bilateral forces are allowed to differ). The required level of bilateral coordination was varied in three conditions (low, medium, high) by varying the projection matrix *c:*

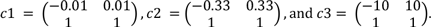

Each projection matrix was used in a block of 20 trials, i.e. the duration of a block was 400 s or about 7 minutes. In addition to the required level of synergy, the total force level was varied to ensure that participants were dependent on visual feedback for successful task performance. The force level necessary during the constant force part varied between two levels (1.7 and 2.1 N per hand) and trials with different force levels were randomized within a block. The order of the 3 blocks was counterbalanced between participants. The inter-trial interval was 5 s during which the performance score was presented (percentage of time the cursor was within the target) and the total duration of each trial was hence 20 s. There was a two-minute break between the blocks.

### 2.3 Data acquisition

Corticomuscular coherence and intermuscular coherence were assessed using electroencephalography (EEG) and high-density surface electromyography (HDsEMG). EEG was recorded using a 64-electrode nylon cap placing the electrodes according to the extended 10-20 system and amplified using a 64-channel Refa amplifier (TMSi, Enschede, The Netherlands; sampling rate 1024 Hz). HDsEMG was recorded using two 64-channel (8×8, IED 4 mm) electrode grids (Fig. 1A) and amplified using a 128-channel Refa amplifier (TMSi, Enschede, The Netherlands; sampling rate 2048 Hz). The electrode grids were placed over the flexor pollicis brevis (FPB) muscle of both hands. Pinch grip force was recorded using a custom-made force sensor (Fig. 1A; Boonstra et al. 2005) and amplified using a National Instruments amplifier (NI SCXI-1121, sample rate 1000 Hz). At the start of each trial, the force sensor amplifier sent a synchronization pulse to both Refa amplifiers, which was used to align data offline.

### 2.4 Preprocessing

The data analysis was performed in Matlab (2012a, The MathWorks, Natwick, MA, USA) using the Fieldtrip toolbox for M/EEG analysis (http://www.ru.nl/neuroimaging/fieldtrip; Oostenveld et al. 2011). EMG, EEG, and force signals were aligned using the aforementioned synchronization pulse and segmented into separate trials. Force signals were linearly interpolated to 2048 Hz. Only the constant force part of each trial was selected to asses bilateral muscle synergies, i.e. from t = 6 to t = 15s (excluding the first second of constant force to avoid transient activity). EEG signals were linearly interpolated to 2048 Hz and high-pass filtered (Butterworth, second order, cut off 1 Hz). The signals were band-stop filtered at 50 Hz and its higher harmonics to remove line noise. ICA was used to remove eye-blinks, eye-movements and muscle activity from the EEG data (Jung et al. 2000).

We used HDsEMG to improve the estimation of corticomuscular and intermuscular coherence (van de Steeg et al. 2014). Bad HDsEMG channels were removed and on average 59±4 channels per hand were used for further analysis. The HDsEMG signals were re-referenced to the reference electrode (placed on the process styloid of the radius), high-pass filtered (Butterworth, second order, cut off 10 Hz) and band-stop filtered at 50 Hz and its higher harmonics to remove line noise. Principal component analysis was applied to the HDsEMG signals to reduce the effects of amplitude cancellation and heterogeneity in the motor unit action potentials and to improve the estimation of correlated input (Staudenmann et al. 2006; van de Steeg et al. 2014). On average, the leading four principal components were removed (left hand 3.6 ± 7.2, right hand 4.5 ± 6.0), and signals were reconstructed using the remaining PCs. After this, the EMG envelopes were extracted using the Hilbert transform. This full-wave rectification provides an instantaneous amplitude estimate of the EMG signal, which reflects the net input to the motor unit pool (Boonstra and Breakspear 2012). EMG envelopes were averaged across all grid electrodes for each hand separately (van de Steeg et al. 2014).

### 2.5 Data Analysis

#### 2.5.1 Muscle synergies

Muscle synergies can be quantified according to the uncontrolled manifold (UCM) method as

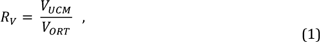

where *V_UCM_* is the variability in the dimension irrelevant for task success (i.e. the UCM) and *V_ORT_* the variability in the dimension orthogonal to the UCM, i.e. the dimension relevant for task success (Latash et al. 2002). Strictly speaking we did not have a completely uncontrolled manifold here, since participants had to control both DOFs in the task parameters (x-and y-position of cursor). Nevertheless, we manipulated the degree with which both DOFs need to be controlled (Nazarpour et al. 2012), and used the UCM method to quantify changes in bimanual coordination. Here we used the variance within a trial as a measure of movement variability (Scholz et al. 2003). This approach is preferable when the number of trials is small. Movement error was computed by subtracting the target forces from the measured force signals. Fluctuations in the UCM and ORT dimensions were estimated by the sum and the difference of the error signals of both hands, respectively. Here the difference of the two forces was the manipulated performance variable and hence could be considered the ORT dimension, i.e. the controlled manifold.

The variances *V_UCM_* and *V_ORT_* were determined within each trial. The ratio of these variances gives the quantitative measure of synergy (*R_v_*) according to equation (1). *R_v_*<1 indicates no synergy is present, whereas *Rv*>1 indicates a synergy between bilateral hand muscles (Latash et al. 2002). To correct for the inherent skewedness of ratio data, the *R_v_* values were log transformed (log *R_v_* > 0 indicates a synergy and log *R_V_* < 0 no synergy) (Verrel 2010). The *R_V_* values were averaged over the 10 trials within each condition.

Previous studies have generally investigated the measure of synergy *R_V_* in the time domain. To investigate the time scale at which the bilateral synergy was established and to investigate the potential relationships with corticomuscular coherence and intermuscular coherence, we extended the measure of synergy *R_V_* to the frequency domain. Scholz et al. (2003) already showed that there seem to be different time scales involved in the structure of movement variability. The power spectral density (PSD) describes the distribution of the variance across frequencies, such that the integral of the PSD across frequencies is equal to variance of a signal. Hence, equivalent to equation (1) the measure of synergy *R_V_* in the frequency domain is given by:

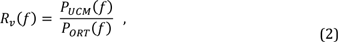

where *P_UCM_*(*f*) and *P_ORT_*(*f*) are the PSD in UCM and ORT dimension, respectively. The PSDs in the UCM and ORT direction were estimated using a multi-taper method based on discrete prolate spheroidal sequences (Slepian sequences) as tapers (Mitra and Pesaran 1999). The amount of smoothing through multi-tapering was set at ± 0.2 Hz. We assessed *R_V_* in the time and frequency domain to compare both approaches.

#### 2.5.2 Corticomuscular and intermuscular coherence

We first localized the sources in the brain showing maximal corticomuscular coherence with the EMG recorded from the left and right hand using DICS beamformers (Gross et al. 2001). Since we did not have access to individual subjects anatomical MRI data, we used the volume conduction model, source model and MRI templates integrated in the Fieldtrip toolbox (Oostenveld et al. 2011). Using this method, first the cross-spectral density (CSD) between all EEG signal pairs and between all EEG signals and the HDsEMG signals were calculated, again using a multi-taper method with a center frequency of 23 Hz and ±7 Hz of spectral smoothing (sensor level analysis showed significant corticomuscular coherence in the 16-30 Hz range). We used the averaged CSD over the z-scored data of all participants to localize two average sources. The coordinates of the sources were converted to the Talairach coordinate system to determine the location labels using the Talairach Client (Lancaster et al. 2000).

For the two locations showing maximal corticomuscular coherence with the EMG envelopes of the right and left hand, we constructed a spatial filter for each participant to reconstruct virtual source signals. The virtual source signals contain a magnitude and a direction and hence yield three signals (in *x*-, *y*-, and *z*-direction) for each solution point. Singular value decomposition was used to select the direction in 3D space that explains most variance, thus leaving us with one source signal per participant per hand. Magnitude squared coherence between the virtual source signals and the averaged HDsEMG signals (corticomuscular coherence) and between the averaged HDsEMG signals of the two hands (intermuscular coherence) were calculated over the constant force part (t = 6-15 s). Again, a multi-taper method was used for spectral decomposition. The amount of smoothing trough multi-tapering was set at ±1.5 Hz, which means a 3 Hz smoothing box was used around each frequency of interest. Based on the grand averages the frequency ranges of interest (i.e. showing significant coherence) were selected for statistical analysis.

### 2.6 Statistical analysis

Statistical analysis was performed in IBM SPSS Statistics Version 21. A 3×2 repeated measures ANOVA with within-subject factors *coordination level* and *force level* served to test for significant differences in *R_V_* in the time domain. Likewise, coherence values in the frequency bands of interest were compared using the same 3#x00D7;2 repeated measure ANOVA. Hence, four 3#x00D7;2 repeated measures ANOVA were performed: *R_V_*, corticomuscular coherence between the left hand and it’s virtual source signal, between the right hand and it’s virtual source signal and intermuscular coherence between both hands. The same 3×2 repeated measures ANOVA was used for each frequency band of interest to test for significant differences in *R_v_(f).* The 95% confidence intervals of the coherence estimates were determined through phase randomization. In addition, Spearman’s rank-order correlation was estimated between *R_V_* and the coherence values across the 6 (3×2) conditions for each participant separately, to directly test the relationship between neural synchrony and muscle synergies. The correlation coefficients of the 16 participants were compared against zero using a one-sample t-test to test the relationship at group level.

## 3. Results

Figure 2 shows force data of a representative participant and demonstrates the effectiveness of the experimental manipulation: In the low coordination condition the two forces were allowed to differ from each other (Fig. 3A and B) and the variability was most pronounced in the ORT dimension (Fig. 3C). In contrast, in the high coordination condition the two forces differed relatively little from each other and variability was mostly observed in the UCM dimension.

**Figure 2.**
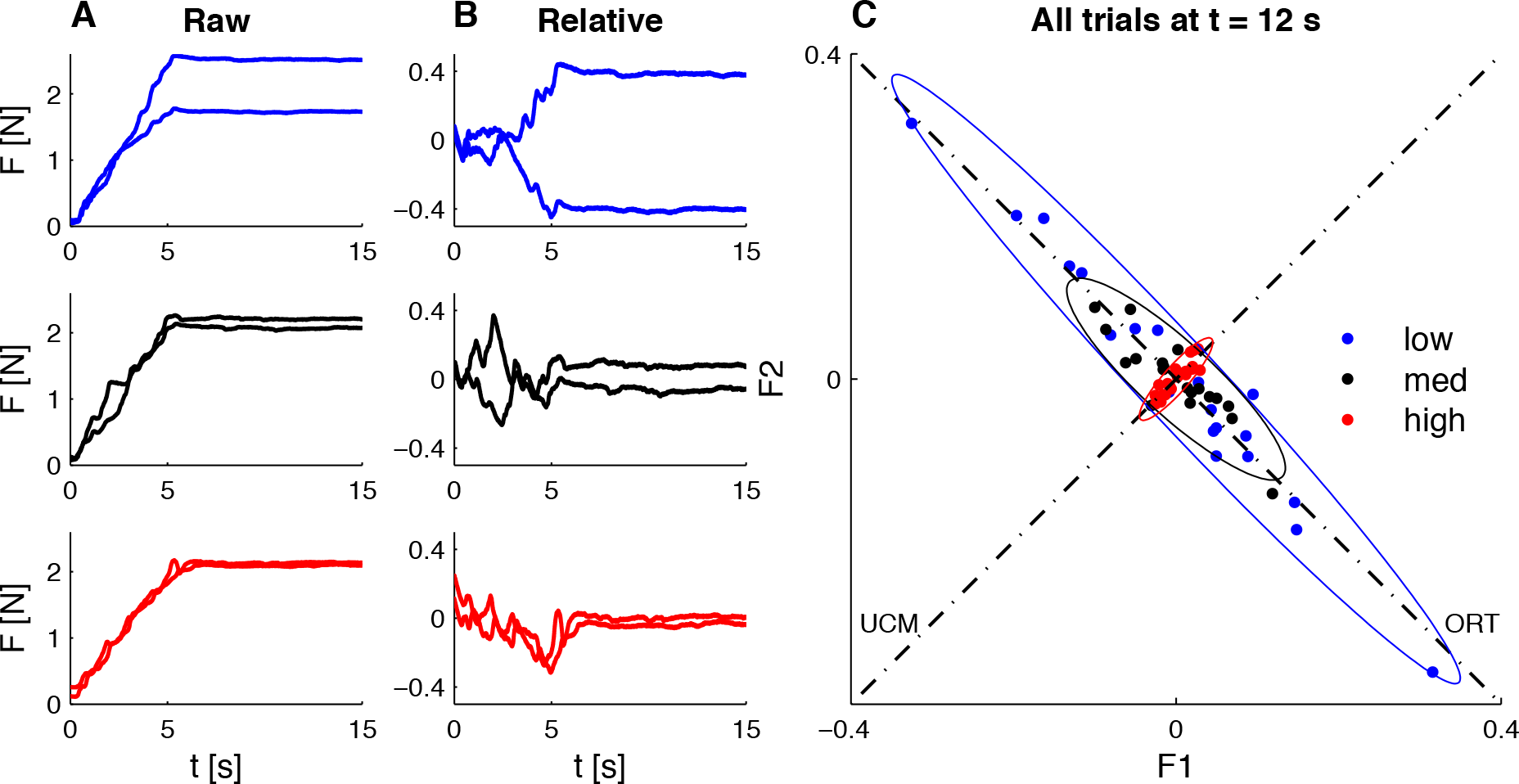
Force data of exemplar participant. The rows show the three coordination levels (blue = low, black = medium, red = high). **A)** Time series of raw force signals of both hands for three representative trials. **B)** Time series of error signals (force signals relative to target signals). **C)** Cluster plot with on the horizontal and vertical axes force levels of the left and right hand, respectively, at *t* = 12 s for all 16 participants. Values in panel C are relative to the average over those trials. Ellipses show the 95% confidence interval.

*R_V_* is calculated according to Equation (1) to quantify the level of muscle synergy for each condition separately (Fig. 3). Log *R_V_* revealed a strongly significant main effect of synergy level (*F*_(2, 30)_ = 182.36, *p* < .001). No significant main effect of absolute force level and no interaction effect were found. As stated, log *R_V_* < 0 and log *R_V_* > 0 quantify no synergy and synergy, respectively. One-sample t-tests revealed that log *R_V_* was significantly smaller than 0 in the low coordination condition for both force levels (p < .001), significantly larger than 0 in the high synergy condition for both force levels (*p* < .001), and not significantly different from 0 in the medium synergy condition for both low (*p* = .15) and high (*p* = .33) absolute force levels. These results confirmed that the experimental manipulation resulted in the intended differences in bimanual synergy between conditions.

**Figure 3.**
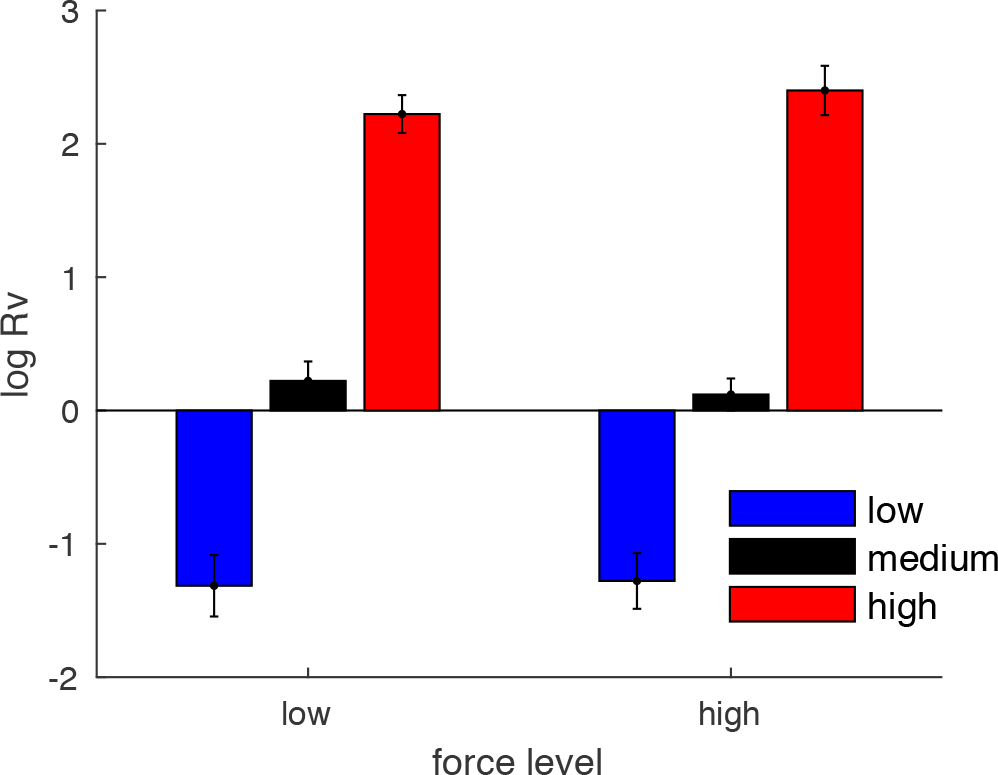
The level of muscle synergy Rv across experimental conditions. Values are averaged over the 10 trials per condition (blue = low, black = medium, and red = high coordination). Error bars show the standard error of the mean.

Figure 4 shows the power in both dimensions and *R_V_* as a function of frequency. As expected, most of the power was observed in the lower frequencies (Fig. 4A, B). For low frequencies (< 0.5Hz) the power in the orthogonal dimension decreased with increasing coordination (high condition). In contrast, the power in the UCM dimension increased with increasing coordination level. Consequently, *R_V_* was higher in the higher coordination conditions for these low frequencies (Fig. 4C). Figure 4D shows a different frequency profile for different coordination conditions: For low and medium *R_V_* was negative for very low frequencies (< 0.5Hz), positive for frequencies between 0.5 and 2 Hz, and then converged to 0 at higher frequencies. The frequency spectrum for the high coordination condition was largely opposite.

The pattern of *R_V_* at frequencies below 0.5 Hz hence largely mirrored the time-domain analysis shown in Figure 3, which is expected as these low frequencies contain most power and thus largely determine the overall variance of the signals. However, the order of *R_V_* across conditions changed at higher frequencies. For frequencies of 0.5 - 2 Hz the largest *R_V_* was observed in the low coordination condition and decreased for the medium and high coordination conditions (Fig. 4D). In this frequency band, the strongest synergy is thus observed when the lowest bimanual coordination is required. In frequencies above 2 Hz the order across conditions changed again and was more similar to the order seen at the lowest frequencies, that is, the strongest synergy was observed in the high coordination condition. ANOVA of *R_V_*(*f*) showed a significant main effect of synergy level for frequencies below 0.5 Hz (*F* (2,30) = 35.96, *p* < 0.001), for frequencies between 0.5 and 2 Hz (*F* (1.4,20.8) = 8.82, *p* = 0.004) and for frequencies above 2 Hz (*F* (1.6,23.8) = 7.47, *p* = 0.005). There was no significant main effect of force level and no significant interaction effect for any of the three frequency ranges.

**Figure 4.**
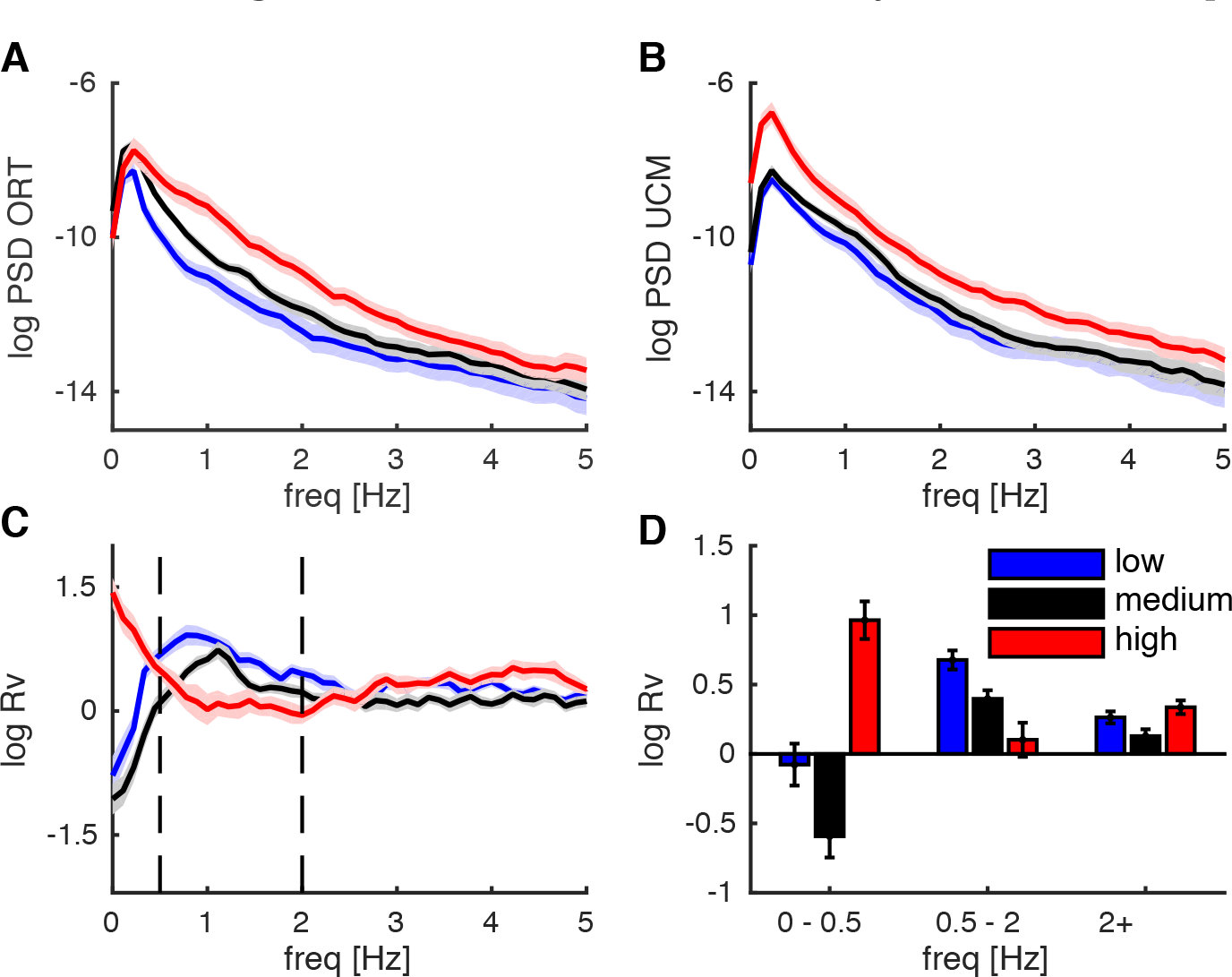
UCM analysis in frequency domain. The solid lines and bars represent grand averages, averaged across the 20 trials per coordination level and all participants. In the figure we averaged over the two absolute force levels within each coordination level. The shaded areas surrounding the lines and the error bars in the bar graph represent the standard error of the mean. Each colour shows one of the three conditions (blue = low, black = medium, red = high). **A)** Power spectral density in the orthogonal dimension (psd ORT) displayed on a logarithmic scale. **B)** Power spectral density in the uncontrolled manifold dimension (psd UCM). **C)** *R_V_* in frequency domain. Vertical dashed line delineate the different frequency bands. **D)** R_v_ divided in three frequency bands, i.e. 0-0.5 Hz, 0.5-2 Hz and 2-10 Hz.

DICS revealed the cortical sources showing maximal corticomuscular coherence in the beta frequency range (23 ± 7Hz) for each hand. Figure 5 shows the sources visualized on a general MRI template (Holmes et al., 1998). The coordinates of the sources in the MNI coordinate system were [4.7, 0.5, 5.5] cm and [-4.2, 0.0, 6.0] cm for the left and right hand, respectively. Both sources were in the precentral gyrus (i.e. Brodmann area 6), which is mainly composed of the premotor cortex.

**Figure 5.**
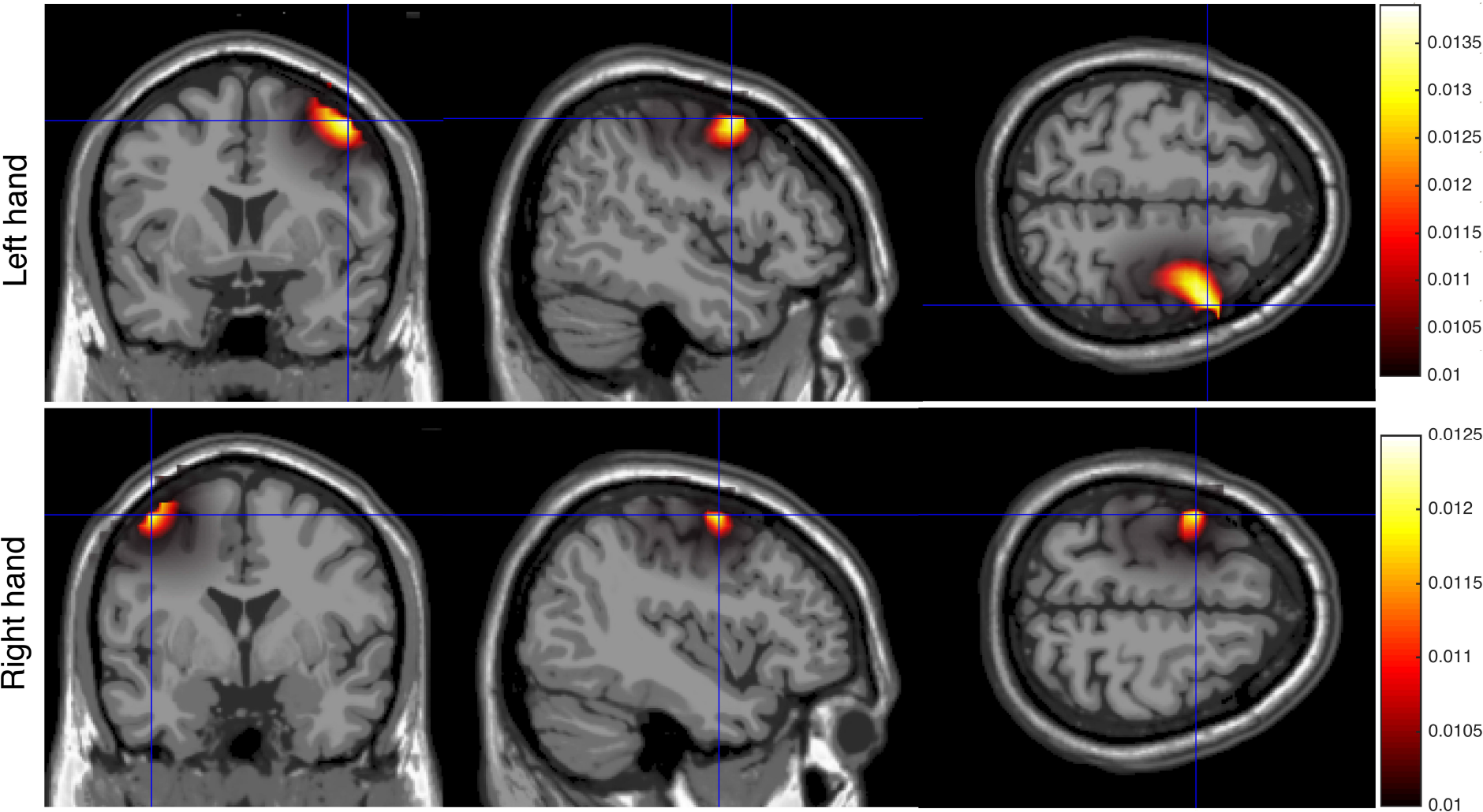
Cortical sources showing maximal corticomuscular coherence. Upper row shows the source with maximal corticomuscular coherence with the EMG envelop of the left hand. Lower row shows the source with maximal corticomuscular coherence with the EMG envelop of the right hand. Orthogonal blue lines intersect at the maximal corticomuscular coherence voxel. Sources are displayed on an MRI template.

As expected, corticomuscular coherence between EMG envelopes and contralateral virtual source signals during the constant force interval was statistically significant in the beta band (16-30 Hz; Fig. 6A). Importantly, there was a significant main effect of condition in corticomuscular coherence between left hand and right cortex (*F*_(2,30)_ = 12.25, *p* < .001) and between right hand and left cortex (*F*_(1.6,23.5)_ = 11.88, *p* = .001). Posthoc comparisons showed higher beta-band coherence in low (*p* = .008) and medium (*p* = .004) compared to high coordination levels between left hand and right cortex, and higher beta-band coherence in low compared to medium (*p* = .01) and high (*p* = .004) coordination levels between right hand and left cortex (Fig. 6C). In contrast, intermuscular coherence was only significant in the alpha-band (5-12 Hz; Fig. 6A) and mainly in the high coordination condition. Intermuscular coherence in the alpha-band showed a significant main effect of condition (*F*_(1.5,22.5)_ = 10.19, *p* = 0.002). Posthoc comparisons revealed larger alpha-band coherence in high compared to low (*p* = 0.015) and medium (*p* = 0.002) coordination levels (Fig. 6C). There was a significant increase in corticomuscular coherence between left hand and right cortex (F(1,15) = 7.09, p = 0.018) and a significant decrease in intermuscular coherence (F (1,15) = 5.15, p = 0.038) at higher force levels. There was no main effect of force level for coherence between right hand and left cortex (F (1,15) = 0.014, p = 0.908), nor were there any significant interaction effects.

**Figure 6.**
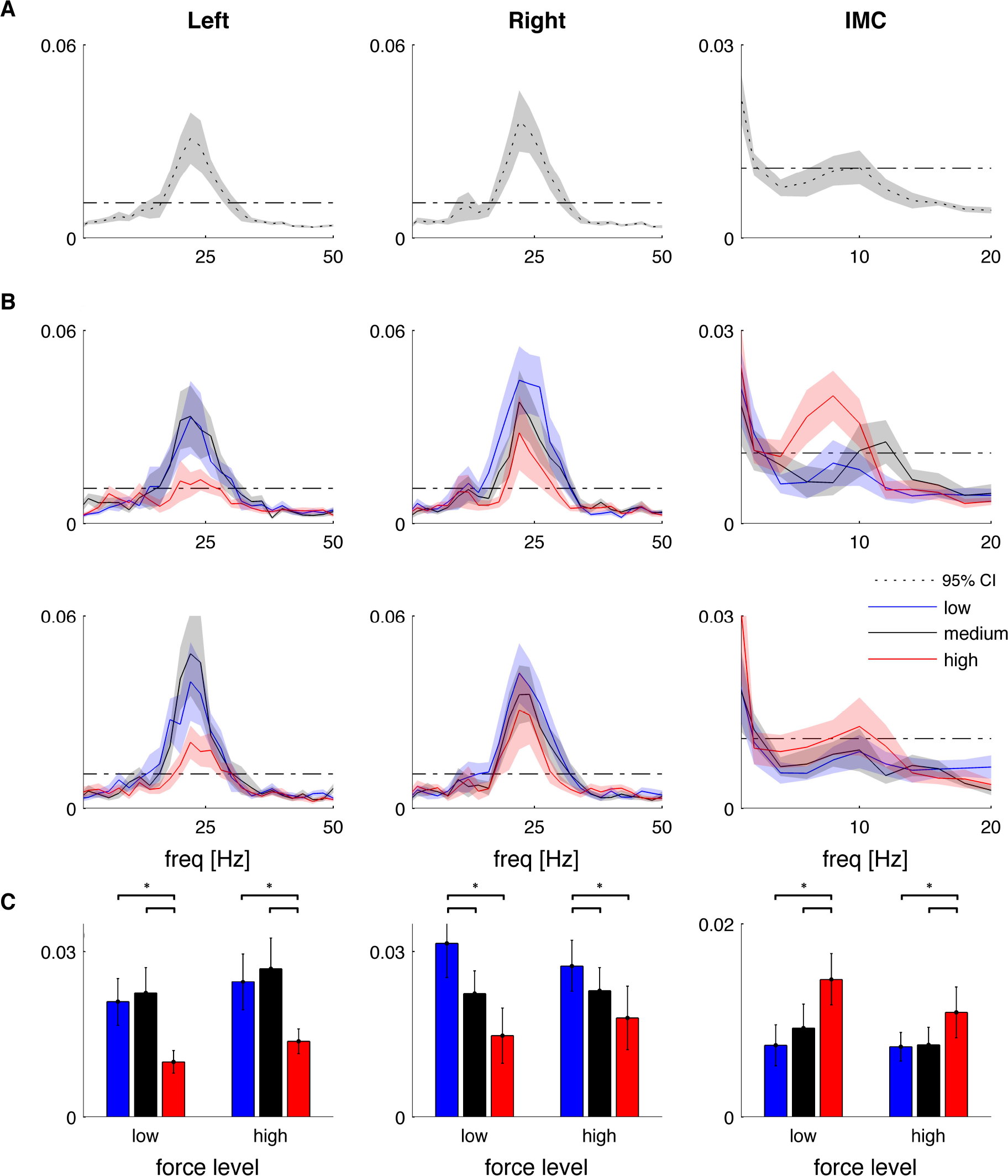
Corticomusuclar and intermuscular coherence. Lines and bars are averages over participants, shaded areas and error bars depict the standard error of the mean and dash-dot lines are the 95% confidence interval. Columns depict from left to right: corticomuscular coherence between left hand and right motor cortex, coherence between right hand and left motor cortex and intermuscular coherence between both hands (IMC). **A)** Grand-average coherence spectra **B)** Coherence in the three coordination conditions (blue = low, black = medium, red = high). The upper and lower row in **B** show low and high absolute force level, respectively. **C)** Averaged coherence in significant frequency bands (16 - 30 Hz for corticomuscular coherence and 5 - 12 Hz for intermuscular coherence). Color-coding is same as in **B**. * significantly different at *p* < 0.05.

When comparing the correlation coefficients of the 16 participants against zero, we found negative correlations between *R_V_* and corticomuscular beta-band coherence with the left EMG (rho = −0.42±0.07, *t*_(15)_ = −5.99, *p* < .001) and right EMG (rho = −0.55±0.09, *t*_(15)_ = −6.00, *p* < .001) and positive correlations between *R_V_* and intermuscular coherence (rho = −0.40±0.10, *t*_(15)_ = 3.90, *p* = .001).

## 4. Discussion

We investigated the relationship between functional connectivity and muscle synergies by experimentally manipulating the level of bimanual coordination. We estimated corticomuscular coherence between the FPB muscle and source activity in the contralateral motor cortex and intermuscular coherence between bilateral FPB muscles at three levels of bimanual coordination. We hypothesized that intermuscular coherence is involved in motor coordination and would hence increase with stronger bimanual synergy levels, while corticomuscular coherence would decrease as it is mainly involved in the control of individual muscles. As expected, corticomuscular coherence in the beta band (16-30 Hz) decreased while intermuscular coherence in the alpha band (5-12 Hz) increased in the condition requiring stronger bimanual coordination. We used the UCM method to quantify the synergy between bilateral hand muscles, which confirmed that the experimental manipulation had the intended effect. We extended this metric to the frequency domain to investigate the time scale at which the bilateral synergy is established. Our analysis revealed distinct frequency bands: for low (<0.5 Hz) and high (>2 Hz) frequencies *R_V_* was highest in the high coordination condition, whereas for frequencies between 0.5 and 2 Hz this effect was largely inversed. When correlating *R_V_* with functional connectivity across conditions, we found negative correlation with corticomuscular coherence and positive correlation with intermuscular coherence. Taken together, our findings suggest that distinct motor pathways are involved in bilateral coordination of hand muscles.

The current study confirms that neural synchronization is a mechanism involved in the formation of muscle synergies by experimentally manipulating the required level of bilateral coordination and showing increased intermuscular coherence with increasing levels of bimanual synergy. These results extend previous studies that have shown a relationship between intermuscular coherence and muscle synergies. Synchronized motor neuron activity is commonly observed in synergistic muscles that act together to accomplish the same joint movement (De Luca and Erim 2002; Laine et al. 2015; Poston et al. 2010). Intermuscular coherence has also been observed between muscle acting on distinct joints, for example between bilateral hand muscles when both fingers moved inphase but not for anti-phase movements (Evans and Baker 2003). Similarly, homologous TA muscles revealed 10-Hz intermuscular coherence when they were simultaneously activated during rhythmic postural sway (Boonstra et al. 2009a). Intermuscular coherence has also been observed for multi-muscle synergies involved in postural control (Boonstra et al. 2015; Danna-Dos-Santos et al. 2014). While these studies are largely observational, we experimentally manipulated the visual feedback to structure the force variability of the left and right hand (cf. Nazarpour et al. 2012). It is important to notice that while we induced changes in correlations between bilateral force fluctuations, the mean force remained unchanged excluding potential effects of force level and amplitude cancellation on intermuscular coherence (Heroux and Gandevia 2013).

The patterns of functional connectivity inform the neural circuitry involved in muscle synergies (Boonstra 2013). Corticomuscular coherence was significant in the beta band (16-30 Hz), while intermuscular coherence was only observed in the alpha band (5-12 Hz; Fig. 6). As bilateral EMG activity in the alpha band was not coherent with cortical activity, it is unlikely that the correlated input generating intermuscular coherence in this frequency band had a cortical origin. This suggests that two distinct neural pathways are involved in corticomuscular and intermuscular coherence (cf. Boonstra et al. 2009b). The two pathways may reflect a phylogenetically newer system containing monosynaptic innervations to motoneurons of individual muscles and a phylogenetically older system containing descending projections driving spinal interneuronal modules, respectively (cf. Bizzi and Cheung 2013). The contribution of both pathways might have been varied depending on the required level of bimanual coordination, which resulted in opposite changes in corticomuscular and intermuscular coherence. By experimentally modifying the degree of unimanual synergies, Nazarpour et al. (2012) found intermuscular coherence in the beta band between upper arm muscles forming a muscle synergy. Intermuscular coherence in the beta band between muscles within the same arm may reflect divergent corticospinal projections (Farmer et al. 1993). Hence correlated input to unimanual muscles may be generated by the phylogenetically newer system. It appears that multiple parallel pathways are involved in the formation of muscle synergies (cf. Tresch and Jarc 2009), rather than a strict hierarchical organization (Loeb et al. 1999; Ting and McKay 2007). The descending pathways (Lemon 2008) and spinal circuitry (Bizzi and Cheung 2013) involve various degrees of divergent projections to distinct motor unit pools. It is hence possible that correlated input observed in muscles forming a synergy results from the activation of dedicated hardwired circuits. However, Nazarpour and colleagues found that muscles that have no direct anatomical connection could rapidly form synergies if required by an abstract new task suggesting that even if low-level anatomical synergies may be used, these can be readily overwritten. The extensive anatomical divergence and convergence in the motor system may thus form a substrate on which abstract task-dependent functional synergies emerge (Nazarpour et al. 2012). Correlated input may result from multiple parallel projections to the muscles and require the coordination of multiple sources that simultaneously constrain muscle activity. Such a many-to-many relationship would involve synchronization throughout the motor system that can be conveniently studied using complex network analysis (Boonstra et al. 2015).

We quantified bilateral synergies by extending the measure of synergy *R_V_* in the frequency domain. These analyses showed that while the low frequencies (0 - 0.5 Hz) mirrored the time-domain analysis, i.e. *R_V_* was highest in the high coordination condition, frequencies between 0.5 and 2 Hz revealed a largely opposite effect. The increased fluctuations in the task-relevant direction *(V_ort_)* observed in this frequency range are somewhat surprising, as it would suggest the strongest synergy is observed in the low coordination condition. The reversal of the order of *R_V_* across conditions indicates that bimanual muscle coordination operates at distinct timescales. This confirms previous findings by Scholz et al. (2003) who also found different timescales involved in the structure of force variability. They suggested control on two hierarchical levels, i.e. high-level voluntary control that reduces the variance in the task-relevant dimension (V_ort_) on a larger timescale (> 250 ms), and low-level involuntary control that channels the variance into the task-irrelevant dimension *(V_ucm_)* on a smaller timescale (< 250 ms). The apparent difference in timescales between our findings and their findings (Scholz et al. 2003) likely reflects their choice of an arbitrary cut-off frequency of 4 Hz (i.e. 250 ms), whereas no cut-off frequency is required when calculating Rv in the frequency domain. Elevated variance at 0.5-2 Hz may reflect an involuntary error correction mechanism, that is, it may result from delays in feedback control. Visually guided responses have a feedback delay of about 200-300 ms, which would correspond to oscillations in force signals with a period length of twice the delay time (i.e. 400-600 ms) when visually tracking a target force (Miall 1996). A visual feedback system may be involved in the high-frequency condition to more closely match bilateral forces, which reduces bilateral variability at longer timescales but increases variability at 0.5-2 Hz. By extending the *R_V_* in the frequency domain the proposed metric enables investigating the timescales of motor coordination in more detail.

Although our findings confirm a relation between intermuscular coherence and muscle synergies, this does not necessitate a causal link and hence does not demonstrate that neural synchrony provides a neural mechanism for establishing muscle synergies. This would require directly perturbing neural synchrony and observing its effect on motor coordination. Instead of being the mechanism for synergy formation, intermuscular coherence may reflect correlated input resulting from diverging motor pathways that are engaged to establish muscle synergies. Intermuscular coherence would then provide the spectral and spatial fingerprints of these pathways and can hence be used to map them (Boonstra 2013; Boonstra et al. 2015). This may help to further establish whether muscle synergies are formed by recruiting hardwired neural circuits or by combing different neural pathways in a task-dependent manner.

## 5. Grants

This research was financially supported by the Netherlands Organisation for Scientific Research (NWO #45110-030).

